# Increase the flow rate and improve hydrogen deuterium exchange mass spectrometry

**DOI:** 10.1101/2022.09.25.509411

**Authors:** Daniele Peterle, David DePice, Thomas E. Wales, John R. Engen

## Abstract

Reversed-phase peptide separation in hydrogen deuterium exchange (HDX) mass spectrometry (MS) must be done with conditions where the back exchange is the slowest possible, the so-called quench conditions of low pH and low temperature. To retain maximum deuterium, separation must also be done as quickly as possible. The low temperature (0 °C) of quench conditions complicates the separation and leads primarily to a reduction in separation quality and an increase in chromatographic backpressure. To improve the separation in HDX MS, one could use a longer gradient, smaller particles, a different separation mechanism (for example, capillary electrophoresis), or multi-dimensional separations such as combining ion mobility separation with reversed-phase separation. Another way to improve separations under HDX MS quench conditions is to use a higher flow rate where separation efficiency at 0 °C is more ideal. Higher flow rates, however, require chromatographic systems (both pumps and fittings) with higher backpressure limits. We tested what improvements could be realized with a commercial UPLC/UHPLC system capable of ~20,000 psi backpressure. We found that a maximum flow rate of 225 μL/min (using a 1×50mm column packed with 1.8 μm particles) was possible and that higher flow rate clearly led to higher peak capacity. HDX MS analysis of both simple and particularly complex samples improved, permitting both shorter separation time, if desired, and providing more deuterium recovery.

## 1. Introduction

Hydrogen-deuterium exchange (HDX) of proteins provides information on conformation, flexibility, interactions, and folding [1]. Rosa & Richards [2] demonstrated that peptide-level information could be achieved by fragmenting a labeled protein – which has incorporated deuterium during solution labeling, usually under physiological conditions (*e.g*. 20-37 °C and pH 7.0-7.5) where protein properties resemble those *in vitro* / *in vivo* – into pieces and analyzing the deuterium level of the pieces. Modern bottom-up HDX measured by mass spectrometry (MS) recapitulates the essential steps of the original experiments: peptides are produced and analyzed under conditions where the exchange is the slowest possible (known as ‘quench conditions’, *i.e*., low pH of 2.5 and low temperature of 0 °C) and then these peptides are separated chromatographically prior to measuring the amount of deuterium [1, 3–5]. The chromatographic step is essential in HDX MS and actually accomplishes at least three things: peptides are resolved from one another before presentation to the mass spectrometer, salts detrimental to ionization are removed, and rapidly exchanging amide hydrogens in amino acid side-chain positions are washed back to hydrogen [4, 6] so the measured deuterium level only reflects incorporation at the protein backbone amide hydrogen positions.

The hydrogen exchange reaction is reversible, meaning deuterium that was incorporated into an analyte during the exchange-in step can exchange back to hydrogen in the H_2_O buffers used in chromatography [1, 4, 5]. It is essential therefore that chromatographic separations in HDX MS be done under quench conditions where this reversion back to hydrogen is the slowest possible, and that separations be completed as quickly as possible; the more label retained, the more information obtained. Fortunately, the pH used for reversed-phase peptide separation is right near the average pH where exchange is the slowest for backbone amide hydrogens, the pH_min_, and thus the optimum quench condition of low pH is met simply by traditional peptide chromatography conditions. Unfortunately, exchange at the pH_min_ is still too fast for high deuterium recovery and therefore, as Rosa & Richards originally showed [2], the temperature is reduced (most commonly to around 0 °C) to maintain the maximum amount of label. Performing a chromatographic separation of peptides at low temperature like 0 °C complicates the separation and leads primarily to a reduction in separation quality (more below). The best HDX MS is ultimately a balance between separation performance and maximum deuterium recovery.

Various parameters post-quench can be optimized to improve the LC/MS steps and concurrently minimize back-exchange and loss of information. One obvious way to improve the separation performance is to use a longer gradient. Even at the low temperature of quench conditions, a long gradient would resolve more species but given more time, more back-exchange and loss of information can occur. Sometimes a longer gradient is used for more complex samples requiring more peak capacity, but unfortunately at the expense of higher deuterium losses. In contrast, small and simple proteins that produce relatively few peptides during digestion can have much faster separations because much lower peak capacity is needed. While temperatures at roughly 0 °C (ice bath) have been the norm since the Rosa and Richards days, subzero separations [7–13] have reported enhanced deuterium recovery due to slower back-exchange. If less deuterium is lost during analysis at −20 °C, for example, the gradient and length of separation could be altered to try to improve the peak capacity and resolving power. One could imagine that if at −20 °C where, for example [7] only 5% of the deuterium is lost, a gradient could be performed that was hours in duration rather than minutes and analyses of extremely complicated samples could be performed. It is clear that improved resolving power is the pathway towards better analysis of very complex samples in HDX MS, something also demonstrated by CE separations for HDX MS [14].

Chromatography of peptides at 0 °C is not optimal. It is well known (*e.g*., [15–19]) that increasing separation temperature decreases solvent viscosity, raises the diffusion coefficient, and leads to a flattened van Deemter curve in the region u > u_p_ (where *u* is velocity and *u*_p_ is the optimal velocity). However, for HDX MS the exact opposite must be done, *i.e*., lowering the temperature. Lowering the temperature of the separation raises solvent viscosity, lowers the diffusion coefficient, and makes the ideal separation more sensitive to flow rate. Some of the effects of lowered temperature can be mitigated by decreasing the particle diameter (with the use of sub 2 μm particles as in UPLC/UHPLC) which has an overall effect similar to increasing the temperature and flattening the van Deemter curve where u > u_p_ [15, 20]. As was previously demonstrated [21, 22], the separation quality of traditional HPLC with 3.5 μm particles at 0 °C is poor whereas separation at 0 °C with 1.7 μm particles using UPLC/UHPLC vastly improves the peak capacity (as predicted by [15]) and separation efficiency for use in HDX MS. Using commercially available UPLC/UHPLC instrumentation, the analysis of seemingly large proteins (> 75kDa) has become routine [1, 23]. Lower chromatographic temperature and smaller particles, however, do come at a price: increased backpressure.

A limiting factor for optimizing chromatographic performance in HDX MS is the recommended backpressure upper limit that a chromatographic system, meaning both pumps and fittings, is capable of handling. As the backpressure of separations at 0 °C is much higher than the backpressure of those same separations at room temperature or higher, chromatographic instrumentation used for HDX MS must have a relatively high backpressure limit to accommodate the required reduction in separation temperature. A general consequence of this high backpressure requirement has been to compromise the ideal flow rate to stay under the backpressure limit of the chromatographic system, unfortunately in the case of HDX MS, to the detriment of the best possible chromatographic efficiency. The introduction of new varieties of UPLC/UHPLC (referred to simply as UPLC from this point forward) pumps and high-pressure fittings offers the potential to improve HDX MS with improved chromatographic efficiency as a result of flowing the mobile phase at flow rates that produce improved separation. Better LC efficiency would greatly enhance HDX MS in terms of the complexity of samples that could be analyzed (for large systems) and the speed with which one could measure peptide deuterium levels (for small proteins).

In this work, we demonstrate the improvements possible in HDX MS separations at 0 °C by flowing much faster (up to 250 μL/min) than is done in many studies (40-65 μL/min). We opted for a practical applied analysis and rather than determining or calculating the theoretical best flow rate for HDX MS at 0 °C, with simultaneous theoretical determination of the optimal particle size and column dimensions which provide the best separation possible at low temperature, we aimed to determine what practical improvements could be realized with a commercially available system that is widely available. We fixed the column geometry to a 1×50mm column packed with 1.8 μm particles, typical for modern HDX MS, and show that a higher flow rate is a possible and obvious improvement. We measured the improvements in terms of peak capacity for protein samples of both low and high complexity. We also compared the deuterium recovery at low and high flow rate.

## 2. Experimental

### 2.1. Reagents

Enolase, alcohol dehydrogenase, bovine serum albumin, and phosphorylase b were purchased from Sigma. More details about these proteins, such as AA length, molecular weight, quaternary structure etc., is provided in Supplemental Fig. S5. Myoglobin was purchased from Sigma-Aldrich (St. Louis, MO, USA). The MassPREP phosphorylase b digestion standard was purchased from Waters (Milford, MA, USA). Pepsin used for protein digestion was from Sigma-Aldrich, part # P6887. Immobilized pepsin resin for digestion was prepared as previously described [24] and packed into an empty 2.1×50 mm stainless steel column (Restek, Bellefonte, PA, USA). Deuterium oxide (D, 99.96%) was from Cambridge Isotope Laboratories (Tewksbury, MA, USA). All other salts, solvents and reagents were of analytical grade and purchased from Sigma-Aldrich or RPI (Mt. Prospect, IL, USA).

### 2.2. Sample preparation

The four-protein mixture were prepared by dissolving lyophilized proteins into 10 mM sodium phosphate buffer, pH 7.4, 150 mM NaCl, 1.75 M GdnHCl. The final monomeric concentration of each protein was 20 μM (5 μM homotetrameric alcohol dehydrogenase, 10 μM dimeric enolase, 20 μM monomeric BSA, 10 μM dimeric phosphorylase b). For phosphorylase b, the protein was reconstituted in 7 M GdnHCl prior to dilution to the final concentration, to facilitate its solubilization. Hydrogen exchange of the 4-protein mixture was performed using 1 μL of protein mixture diluted with 18 μL of labeling buffer (10 mM sodium phosphate buffer, 150 mM NaCl, 99.9 % D_2_O, pD 7.4) and incubated for 10 minutes at room temperature. An undeuterated control was prepared in identical manner except the labeling buffer contained H_2_O rather than D_2_O. After labeling, 19 μL of ice-cold quenching buffer (4 M GdnHCl, 200 mM sodium phosphate, 0.72 M TCEP, pH 2.4) was added before LC/MS.

The myoglobin deuterium recovery experiment was carried out using 20 μM myoglobin resuspended in 10 mM sodium phosphate buffer, pH 7.4, 150 mM NaCl, H_2_O. Hydrogen exchange was performed as described above for the 4-protein mixture. Exchange was quenched with ice-cold 150 mM potassium phosphate, pH 2.4. Maximally deuterated samples of myoglobin were prepared following our previously published protocol [25]. Briefly, 15 μL of myoglobin (10 μM) were lyophilized and resuspended in 7M GdnHCl. The solution was heated at 90 °C for 5 minutes then cooled to 20 °C for 2 minutes. 1 μL of the denatured protein was then diluted with 18 μL of deuterated phosphate buffer, pH 7.4 (same as above) and the exchange was allowed to proceed at 50 °C for 10 minutes. The sample was then cooled at 0 °C for 2 minutes before being acid quenched with ice-cold quenching buffer (as above) and immediately analyzed by LC/MS. Myoglobin labeling experiments were performed in triplicate.

### 2.3. LC/MS

The back-pressure measurements in Fig. 1C were performed using either a Waters nanoACQUITY, an M-Class, or and I-Class UPLC system. The four different ACQUITY UPLC columns (Waters) included two different column geometries (1×50 mm or 1×100 mm) packed with two different stationary phases (BEH 1.7 μM or HSS T3 1.8 μM). Separation efficiency, expressed as peak capacity, was determined by injecting the Waters MassPREP phosphorylase b digestion standard, prepared by dissolving 1 nanomole freeze-dried peptides in 1 mL H_2_O, 0.1% FA. Peptides were trapped and desalted on a VanGuard precolumn trap (2.1×5 mm, Acquity UPLC BEH C18, 3.5 μm) for 3 min at 100 μL/min and eluted with a Waters ACQUITY UPLC HSS T3, 100 Å pore size, 1.8 μm particle size, 1×50mm column by applying different gradient lengths (5-35% acetonitrile in 2, 3, 4, 5, 6, 8, 10 minutes) and flow rates (75, 100, 150, 200, 225 μL/min) as shown in Fig. 2A. For each chromatographic condition, 8 pmols of peptide mixture were injected. Each condition was tested in triplicate.

**Figure 1.**
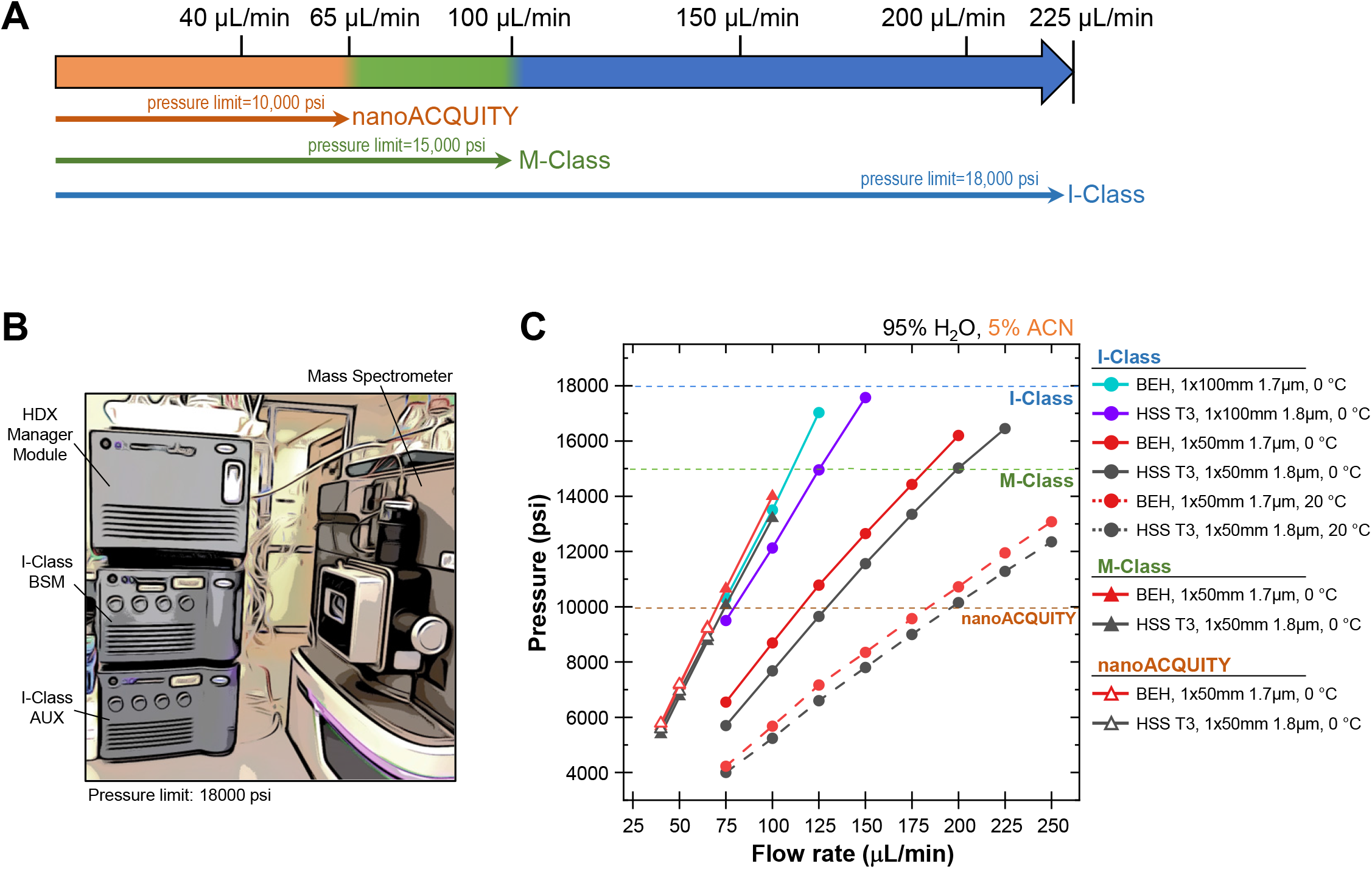
Flow rate and backpressure at 0 °C. (A). Flow rates typically used for HDX MS experiments performed with some of the most common HDX MS platforms (UPLC systems from the Waters Corporation) in order to stay under the backpressure limitations dictated by both the pumps and the fittings. (B). Overall layout of a Waters I-Class UPLC system configured for HDX MS. (C). How flow rate affects backpressure at 0 °C. Measurements of pressure versus flow rate as the UPLC system, column chemistry, column dimensions, and separation temperature were altered. Dotted lines indicate the manufacturer recommended upper pressure limit for each system. In the legend, the column geometry is indicated: stationary phase, IDxlength(in mm) particle size(in μm). These backpressure readings were recorded at 5% acetonitrile, 95% water, 0.1% formic acid; for other solvent compositions, see Supplemental Fig. S1.

**Figure 2.**
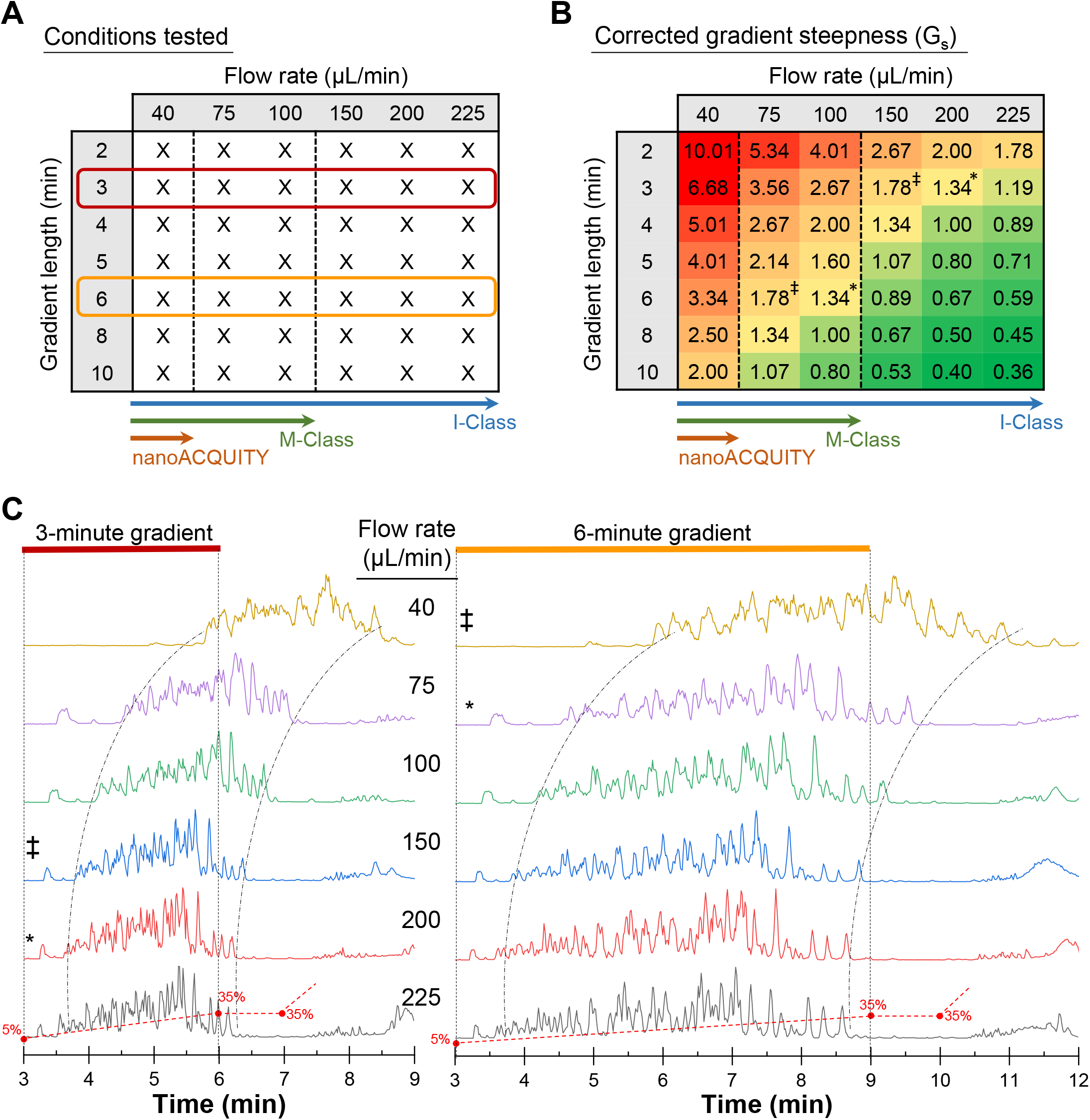
Chromatographic performance at various flow rates. (A). A table showing the flow rates and solvent gradient lengths that were tested and used to calculate peak capacity (Fig. 3). Each condition was analyzed in triplicate for a total of 126 chromatograms. The red and orange boxes highlight the data that are shown as typical examples of chromatography in panel C. Dotted lines indicate the upper boundary for flow rates at 0 °C (see also Fig. 1). (B). Calculation of G_s_ (corrected gradient steepness, expressed as %B per column volume of mobile phase [26, 27]) for the same flow rate and gradient length combinations as in panel A, color coded from worst (red) to best (green) G_s_ based on an optimal value of G_s_ = 0.25-0.33% for reversed-phase peptide separation [33]. A 1×50 mm column with fully porous particles (V_m_ =26.7 μL) and a gradient change of 30% were used for these calculations. See also Supplemental Fig. S2. (C). Chromatograms (Y-axis is base peak intensity) of the 0 °C separation of a phosphorylase b tryptic peptide digest (8 pmol injection, MassPREP standard, Waters) on a Waters HSS T3 column (1×50mm, 1.8 μm particles, 100 Å pores). The gradient was 5-35% acetonitrile (in water with 0.1% formic acid) in 3 minutes (left) or 6 minutes (right) at the flow rates (40-225 μL/min, top to bottom) indicated. Base-peak intensity (BPI) has been plotted. The * and ‡ indicate parameter combinations where the G_s_ values are the same, see panel B. Curved dotted lines are a visual aid to align the gradients.

Chromatography of the four-protein mixture and myoglobin were performed with the same parameters as above, using different gradient lengths and flow rates, but digesting them online using a home-made pepsin column just prior to the trapping/desalting step. For measurements involving hydrogen exchange, the I-Class UPLC was connected to a Waters HDX module (Peltier-cooled chamber) set at 0 °C (temperature is measured in multiple locations on the cooled chamber block, as described in Ref. [22]) and digestion was performed online at 15 °C.

All mass spectrometry was performed with a Waters Synapt XS mass spectrometer using ESI(+) resolution and HDMS^E^ mode. Instrument settings were: 2.5kV cone and 35 V capillary voltages with source and desolvation temperatures of 80 and 175 °C, respectively, and a desolvation gas flow of 600 L/hour. All mass spectra were acquired using a 0.4 sec scan time. The measured peak width averaged 0.11 min at 40 μL/min and 0.03 min for 225 μL/min (see Supplementary Datafile for actual values). Depending on the flow rate, there were between 5.2-17.5 data points per peak, e.g. at 100 μL/min, 6 minute gradient each peak was defined by an average of 9.6 datapoints.

### 2.4. Data analysis

Corrected gradient steepness (G_s_) was calculated as described in [26, 27] using Eq. 1:

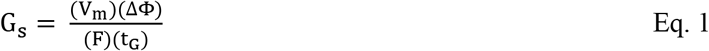

where V_m_ is the interstitial volume (*i.e*., the volume of mobile phase) within the column [calculated using V_m_ = πr^2^LW where r is the column radius (in millimeters), L is the column length (in millimeters), W is the column percent interstitial porosity (0.68 was used for fully porous materials)], ΔΦ is the gradient range, or percent change in organic modifier over the gradient (a 5–35% B gradient is represented as 30), F is eluent flow rate (in milliliters per minute), and t_G_ is the gradient time (in minutes). The V_m_ for the column used (a Waters HSS T3, 100 Å pore size, 1.8 μm particle size, 1×50mm column) was 26.70 μL.

To estimate the sample peak capacity 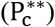, Eq. 2 (according to Dolan, Snyder et al. [28], reviewed in [19]) was used:

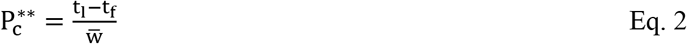

where t_l_ is the retention time for the last eluting peptide, t_f_ is the retention time for the first eluting peptide, and 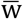 is the average peak width across the separation. While the application of this accepted equation for 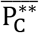 does not produce a precise peak capacity calculation [19, 29], it is suitable for the purposes here in comparing peak capacity while changing various chromatographic variables. Details of the measured peak widths in various separation conditions are found in the Supplemental Fig. S3 and the Supplemental Datafile.

Myoglobin peptic peptides were identified using ProteinLynx Global Server (PLGS) 3.0.1 (Waters) and deuterium incorporation determined using DynamX 3.0 (Waters) as described elsewhere [25]. Deuterium recovery for each peptide, expressed as a percentage, was calculated from the ratio between the experimental deuterium incorporation (Da), corrected for the %D_2_O used (90%), and the theoretical maximum uptake (Da), given by Eq. 3:

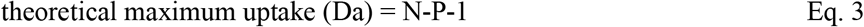

where N is the number of amino acids in a given peptide and P is the number of prolines. If a proline is present at the N terminus, the equation becomes RFU = D / (N-P).

## 3. Results and Discussion

### 3.1. Backpressure measurements

The combination of sub-2 μm diameter stationary phase particles with the 0 °C chromatography temperature required by HDX MS leads to higher backpressure. In order to stay under the backpressure limitation of a chromatographic system, suboptimal flow rates are used, compromising chromatographic efficiency. Improved HDX MS separations should come from using commercial instrumentation with the highest backpressure limits where higher flow rates are possible. Current [30] major commercial UPLC/UHPLC pump vendors and their system backpressure limits include ThermoScientific (Vanquish) ~21 kpsi; Shimadzu (Nexera) ~19 kpsi; Agilent (Infinity II) ~18 kpsi; Waters (I-Class) ~18 kpsi.

A large percentage of HDX MS currently performed uses Waters UPLC systems, including our own research, and our studies were therefore specific to this very common HDX MS chromatographic platform. The latest generation of Waters UPLC pumps, I-Class pumps, have a backpressure limit of ~18,000 psi (1241 bar), and even without considering HDX MS, are an improvement over the original Waters nanoACQUITY or the Waters M-Class UPLC pumps. Unlike previous generations of Waters UPLC pumps, I-Class pumps were not engineered to flow at very low (nanoflow) rates; hence, the pressure and flow sensors necessary for accurate delivery of solvents at sub-microliter flow rates were eliminated from the plumbing in I-Class systems, relative to the M-Class and nanoACQUITY series. The expectation, therefore, is that HDX MS should improve with I-Class systems because higher flow rates that do not exceed the backpressure limit are likely possible, all while still maintaining the 0 °C conditions needed for HDX MS.

Figure 1A summarizes the flow rate ranges at 0 °C that do not exceed the backpressure limits (both of the pumps and the fittings) for three Waters UPLC systems. While acceptable flow rate ranges for both the nanoACQUITY system and the M-Class system were already known [5, 22], the 0 °C flow rate range of the I-Class that did not exceed the backpressure limit had to be determined. We measured backpressure in a dual pump I-Class system coupled to an HDX manager configured for HDX MS (Fig. 1B, Fig S1A) using typical HDX MS experimental conditions, varying the flow rate, column dimensions, and column chemistry. All measurements were performed at 0 °C except for two comparison measurements for the I-Class system at 20 °C. We compared the backpressure readings in the I-Class system to backpressure measurements from both an M-Class system and a nanoACQUITY system configured for HDX MS (Fig. 1B). Standard recommended fittings, which could withstand recommended backpressure limits (~18,000 psi for I-Class; 15,000 psi for M-Class; and ~10,000 psi for nanoACQUITY), were used in all cases. The results (Fig. 1C, Fig. S1B,C) show that with an I-Class system, 0 °C separations can be performed, without exceeding the recommended backpressure limitations at 5% acetonitrile, at flow rates up to 225 μL/min for a 1×50 mm column with 1.8 μm particles and up to 125 μL/min for a 1×100 mm column with 1.8 μm particles. In contrast, the maximum flow rate – without exceeding the recommended backpressure (1 × 50mm column with 1.8 μm particles) – in 0 °C separations with the M-class was 100 μL/min and with the nanoACQUITY system it was 65 μL/min. In summary, the higher backpressure limit of the Waters I-Class UPLC system allows for more than double the flow rates under HDX MS conditions than those afforded by previous generation UPLC systems. The concept can be extended to other UHPLC vendors as well with the general guideline that one should fully exploit the capability of the pumping system being used to flow as fast as the system pressure limit will allow while maintaining 0 °C quench conditions.

### 3.2. Flow rate and gradient steepness

For peptide separations at room temperature and above, it has been shown both theoretically and practically that higher flow rate and longer gradient length concomitantly improve peak capacity for peptide separations [19, 31, 32]. While none of the previous studies were at 0 °C, the trends are expected to remain, although the overall peak capacity should be numerically lower at lower temperature [32] for the flow rates in question (up to 250 μL/min). We systematically investigated peptide chromatography at 0 °C in an I-Class HDX MS system using a series of flow rates ranging from 40-225 μL/min and gradient lengths from 2-10 minutes (Fig. 2A). Chromatograms (mass spectrometry used for detection) of the separation of phosphorylase b tryptic peptides, for which there were at least 76-180 species (assuming 0-1 missed cleavages in theoretical trypsin digestion of UniProt P00489), were obtained in triplicate for all 42 flow rate and gradient length combinations using a 1×50mm HSS T3 column packed with 1.8 μm particles, 100 Å pores. Before interpretation of the spectra we calculated the corrected gradient steepness (G_s_) of each flow rate:gradient combination (see also Fig. S2) to predict the performance of each combination. A good G_s_ range for high-quality separations of peptide mixtures with reversed-phase chromatography at room temperature is 0.25-0.33%B per column volume [33]. With these values in mind, we color-coded the calculated G_s_ for our tested flow rate:gradient combinations (Fig. 2B) and not surprisingly, faster flow rates with longer gradients are predicted to be much closer to the ideal range of 0.25-0.33%B per column volume and should provide better separation efficiency. For slower flow rates, such as 40 μL/min or 75 μL/min, the G_s_ was 6-30 or 3-15 times higher than 0.33, respectively; the gradient changed much too fast for the flow rate.

The theoretical predictions based on G_s_ were then compared to measured chromatographic results at 0 °C. Fig. 2C shows example chromatograms for the 3- and 6-minute gradients at each of the tested flow rates (all chromatograms shown in Supplemental Fig. S2C). Regardless of the gradient length, separation was visibly improved at higher flow rates, with sharper peaks clearly evident. Retention time remained relatively constant regardless of flow rate or gradient (Fig. S3A) but the peak width decreased with higher flow rate (Fig 2C, Fig. S3B,C), consistent with a reduction in longitudinal diffusion and a G_s_ much more appropriate for the column volumes of mobile phase passing through the column. The clear conclusion without going any further is that for peptide separations at 0 °C, higher flow rates provide better chromatographic performance.

### 3.3. Peak capacity measurements

To quantify the chromatographic results (Fig. 2C), the sample peak capacity, 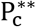, (according to Dolan, Snyder et al. [28], reviewed in [19]) was calculated using Eq. 2. Sample peak capacity is a more useful value to consider for complex samples [32] rather than peak capacity. We determined the retention times of the first- and last-eluting species from the phosphorylase b tryptic digest separations, as well as the average peak width across the separations (see Supplemental Fig. S3 and Supplemental Datafile). The results are shown in Fig. 3. The peak capacity determinations from these measurements ranged from ~5.4-60, which not surprisingly, is lower than peak capacity in the low 100s predicted for peptide separations with longer gradients (say 10-15 minutes) at room temperature [19, 32]. Higher flow rates produced the best peak capacity at 0 °C.

**Figure 3.**
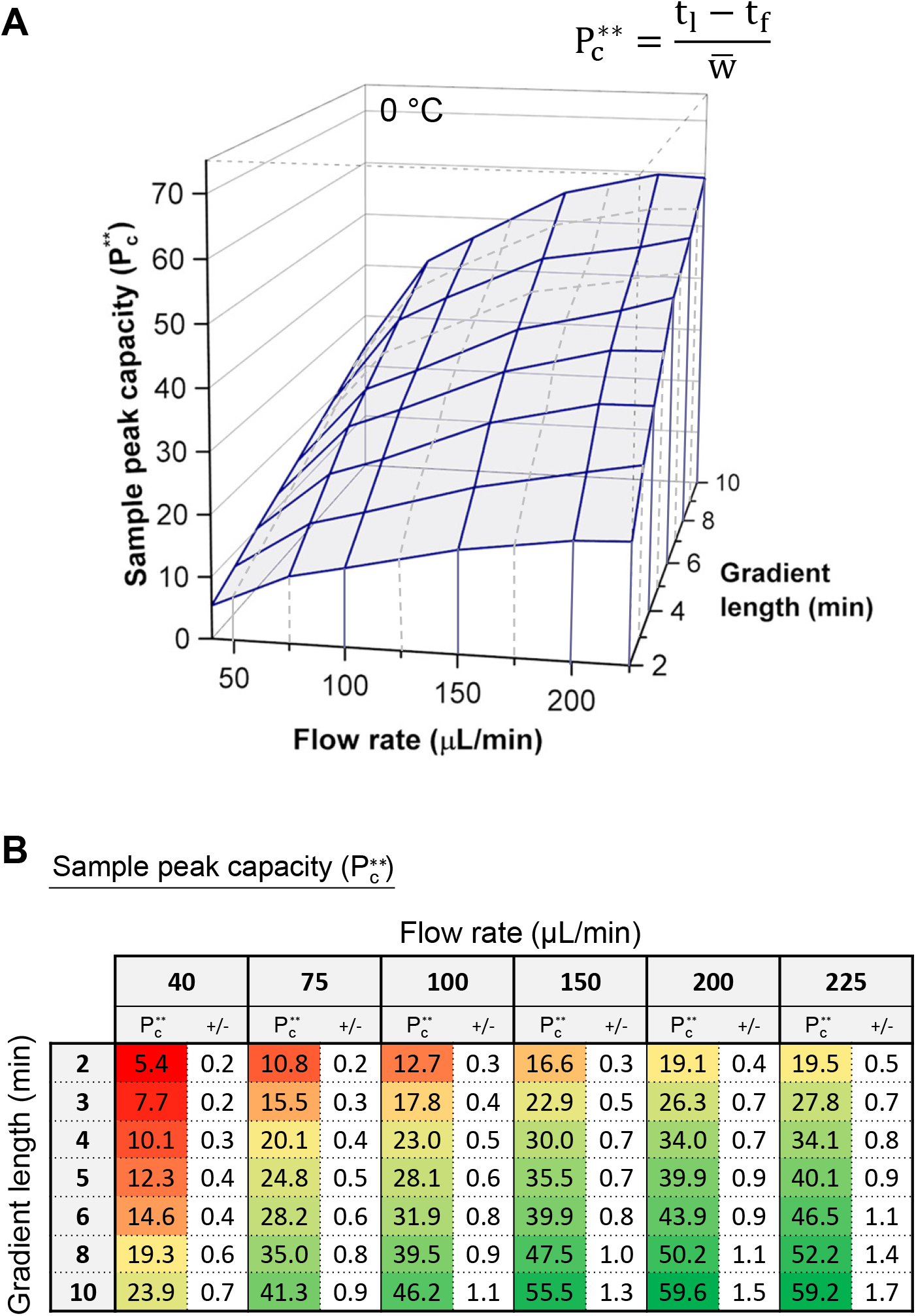
Peak capacity for 0 °C peptide separations. (A) Sample peak capacity 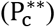 was calculated for the flow rate and gradient length combinations in Fig. 2A using Equation 2 (shown; [28] reviewed in [19]) where t_l_ and t_f_ are the elution times of the last and first peptide, respectively, and w is the average peak width across the 100 most intense peptides identified by PLGS. See Supplemental Fig. S3 and Supplemental Datafile for the measured values and calculations. (B) Numeric values of all calculated sample peak capacities for all tested conditions in panel A, colored from worst (red) to best (green). The uncertainty shown (+/−) is the confidence interval at 95% confidence, calculated as described in Fig. S3 with the actual values provided in the Supplemental Datafile. The best peak capacity for peptide separations at 0 °C is obtained at the highest flow rate and the longest gradient. See also Fig. S4.

An average HDX MS practitioner 10 years ago may have routinely used 40 μL/min as the standard flow rate and these results show how inferior that is compared to what is possible with systems that can accommodate a higher flow rate. For a relative comparison, we calculated the relative percent improvement from a single base condition (Fig. S4). As an example, we selected the base condition of a 6 minute gradient at 100 μL/min, something typical of an M-class separation one might find in recent publications (e.g., Refs. [5, 34]). There is a 54% improvement by flowing at 100 μL/min rather than 40 μL/min, and a further 46% improvement by flowing at 225 μL/min rather than 100 μL/min, which when combined produces a 100% improvement by flowing at 225 μL/min instead of 40 μL/min. Switching from a 6 minute gradient to a 3 minute gradient but holding the flow rate constant at 100 μL/min reduced the peak capacity by 44%. A peak capacity improvement of 86% could be realized by changing from 100 μL/min 6 minute gradient to a 225 μL/min 10 min gradient. One could estimate how much their separation efficiency might decrease/increase using any starting condition as the reference point.

### 3.4. A complex peptide mixture

To study the 0 °C separation of a very complex mixture, something that has many more peptides than the phosphorylase b tryptic digest, we prepared a four-protein mixture to use as a surrogate for a large protein complex. A mixture of four proteins (Fig. S5) that together total 501 kDa in mass quantity and 2,208 amino acids of unique sequence [35], was made, similar to the mixture used previously for early characterization of UPLC in HDX MS [22]. A fixed gradient time of 10 minutes was chosen and the separation monitored as the flow rate was varied from 40-200 μL/min. The chromatographic separations are shown in Fig. 4. As was the case for a simpler mixture, faster flow rate is clearly producing a better separation indicated by sharper peaks (see width measurements in Fig. S6). Considering the mass spectra (Fig. 4B) and selecting a field of ions at any chromatographic point, as chromatographic separation improves so too does the simplicity of ions seen per scan – scan time here is 0.4 sec. Separation of digests of this mixture at 200 μL/min resulted in cleaner m/z spectra versus 100 μL/min. With fewer ions in a given m/z field, it will be more straightforward for software algorithms and manual data processing to identify and characterize isotope windows for deuterated species, an effect that ultimately translates into improved experiments for very large and complex samples where peak overlap may begin to hinder data processing.

**Figure 4.**
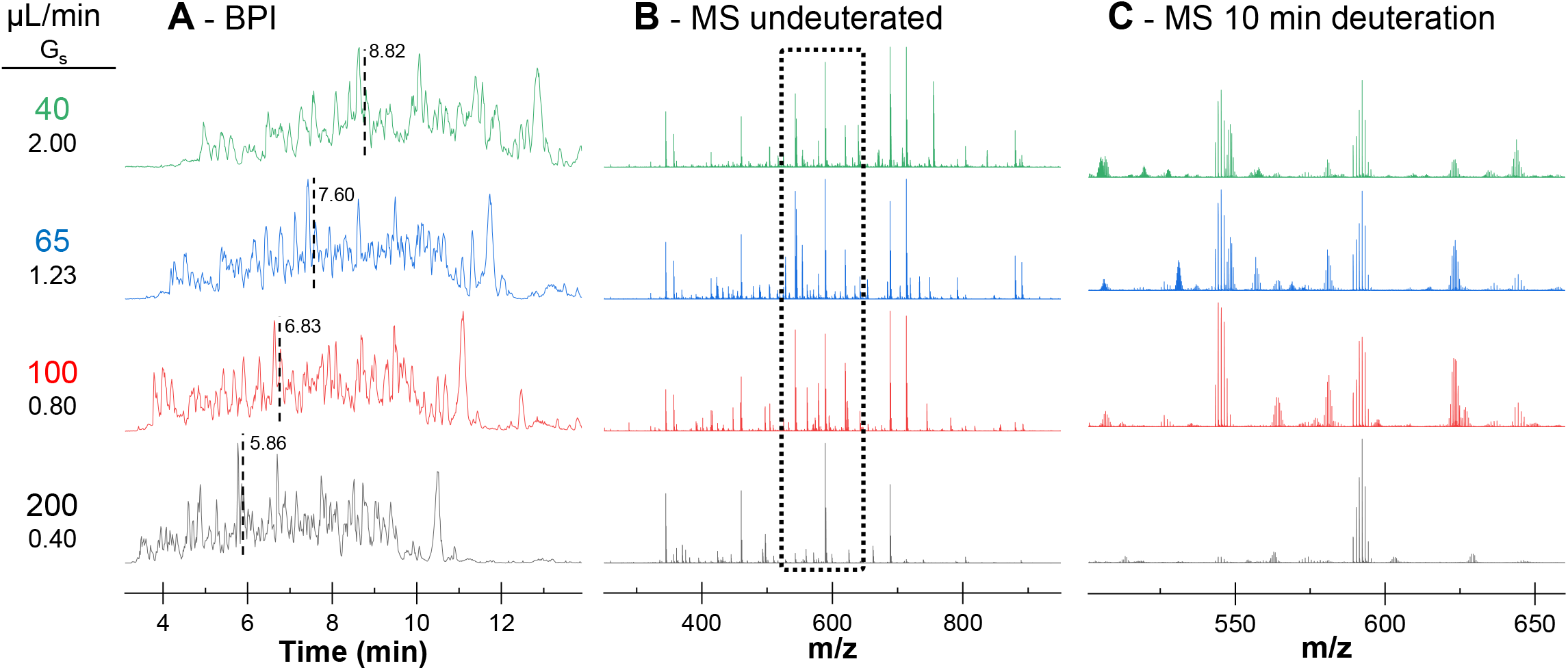
Chromatographic performance with a complicated peptide digest. (A) Chromatograms (BPI on Y axis) of the separation of the four-protein mixture (see Supplemental Fig. S5) during a 10 minute 5-35% acetonitrile gradient, using the same column as described in Fig. 2. The flow rate is shown at the left of each chromatogram along with the G_s_ value for each gradient length and flow rate combination (see also Supplemental Figs. S2, S6). (B). Mass spectra of an undeuterated sample at the times indicated in panel A (vertical lines) where the field of ions is nearly the same. (C) Zoom of the region m/z 500-650 (dotted box in panel B), but with a sample that had been deuterated for 10 minutes and analyzed under standard HDX MS quench conditions.

### 3.5. Deuterium recovery at higher flow rate

Could a change in flow rate from 40 to 200 μL/min induce frictional heating and thereby warm the separation column enough (even for the short amount of time deuterated peptides were exposed to the column) to affect back exchange in HDX MS? If there were more back exchange, and therefore loss of deuterium, during analysis at higher flow rate, this would be undesirable. While a flow of 200 μL/min is faster than used in the past for HDX MS, we hypothesized that it is likely not swift enough to become an issue. In the literature, frictional heating is detectable with larger diameter columns and flow rates much higher (more like 1 mL/min) than those used in our work. Colon et al. [36] showed that 1 mm columns packed with 1.5 μm particles did not generate significant heating power and there was no deterioration of separation efficiency at pressures up to 20,000 psi; the same was found for 2.1 mm columns with 1.7 μm particles at linear velocities of 2.5 mm sec^−1^ (~520 μL/min). Gritti et al. [37] found that column heating occurs at flow rates of >600 μL/min using a 2.1×50 mm BEH column. In the present HDX MS work we used a maximum of 225 μL/min with a 1×50mm BEH/T3 column (1.7-1.8 μm particles), approximately 3 times slower than the slowest rate reported by Gritti et al. and 2 times slower than Colon et al. where almost no heating was observed. Makarov et al. [38] monitored the temperature along the length of BEH 2.1×50 mm 1.7 μm columns at various flow rates from 800-1200 μL/min, with water, methanol and acetonitrile as the mobile phase. They observed heating of 3-5 °C at 800 μL/min in pure water while in pure acetonitrile, heating was cut nearly in half. During an acetonitrile gradient, potential heating will be less of a factor as the gradient progresses, as the acetonitrile (%B) value increases. It is not straightforward to compare our conditions to those of the Makarov et al. studies: the maximum flow rate of our HDX MS measurements (225 μL/min) was 3.5 times less than the slowest flow rate (800 μL/min) monitored in the Makarov et al. study, and our column diameter was 1 mm not 2.1 mm. In addition, our column was actively cooled in a 0 °C chamber while their columns were not. One predicts that frictional heat should be readily dissipated from a 1 mm diameter column [39] especially one maintained in a 0 °C chamber. In conclusion, published measurements suggest that at the flow rates compatible with 0 °C separation for HDX MS (225 μL/min and less), frictional heating will be very small to negligible for sub 2 μm particles in column geometries of 1 mm diameter and probably also of 2.1 mm diameter.

Given deuterium recovery is sensitive to the temperature, we were in a position to measure the effect that higher flow rate had on deuterium recovery and directly experimentally address the question of did higher flow rates lead to frictional heating and higher column temperature that translated to additional loss of deuterium during separation. We measured both sequence coverage and back exchange in 100 μL/min, 6 min gradient at 0 °C and a 200 μL/min, 3 min gradient at 0 °C. The G_s_ for these two conditions was identical (see Fig. 2B) and the 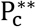 for 200 μL/min, 3 min conditions using the phosphorylase b test mixture was found to be slightly higher than the 100 μL/min, 6 min conditions (see Fig. 3B). The coverage maps for the two conditions were essentially identical (Fig. S7). Given the simplicity of a myoglobin pepsin digestion, a 3 minute separation is more than adequate to resolve all peptides of interest. Deuterium recovery was on average 3.6% higher (range 0.5-9.7%) with the 200 μL/min 3 min gradient (Fig. 5), most likely a result of the faster separation and exposure to back exchange conditions for only half the time of the 100 μL/min, 6 min conditions. We conclude based on both theory and measurement that frictional heating is practically inconsequential for accelerating back exchange in HDX MS at flow rates of 200 μL/min at 0 °C. Even if there is a small amount of frictional heating, it is kept mostly in check by the cooled chamber and 1 mm column diameter and there is an overall improvement in deuterium recovery at higher flow rate compared to the lower flow rates currently used by many in HDX MS (40-65 μL/min).

**Figure 5.**
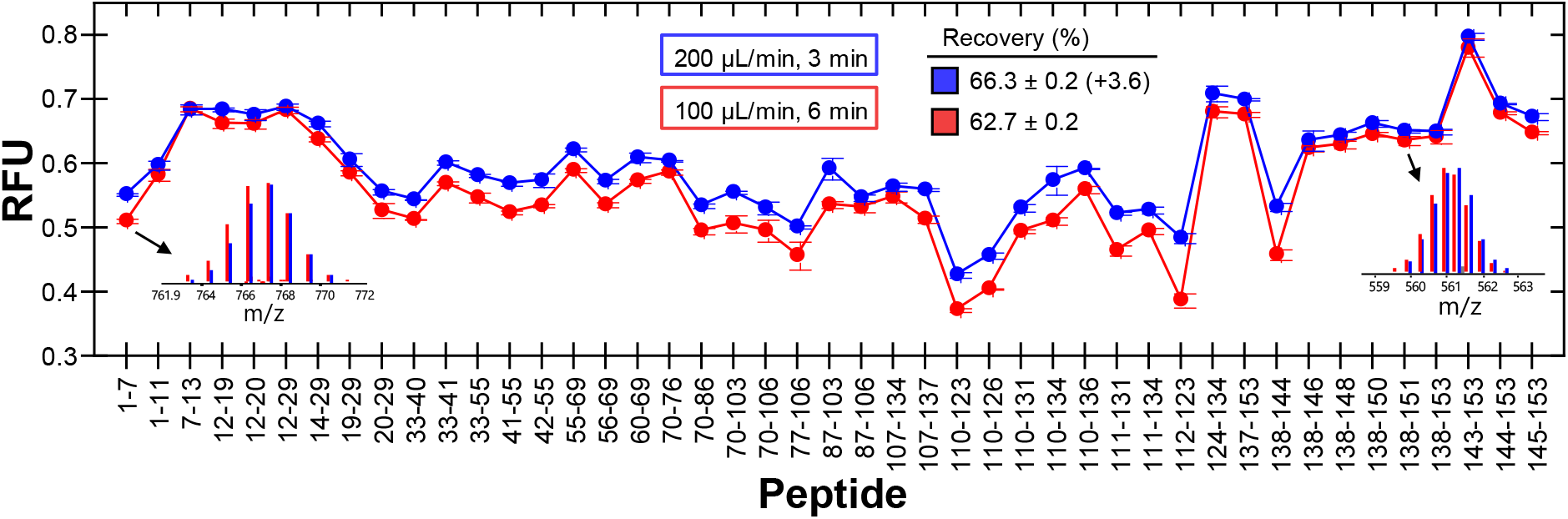
Deuterium recovery at different flow rates. (A) Deuterium recovery was measured using standard HDX MS conditions for a maximally deuterated sample of myoglobin (described in [25]) at 100 μL/min, 6 min gradient (red) and 200 μL/min, 3 min gradient (blue) for the peptic peptides shown. Relative fraction uptake (RFU, y-axis) is shown for each peptic peptide monitored (x-axis). Each datapoint is the average of triplicate measurements and the error bar shown is the error calculated by the DynamX software. The ± range in the legend is the 95% confidence interval. (Insets) Representative mass spectra for two myoglobin peptic peptides comparing overlaid mass spectra for both flow rates.

## 4. Conclusions

We have shown that faster flow rate means improved chromatographic performance for HDX MS. Higher flow rate leads to both higher peak capacity while permitting both shorter separation time and more deuterium recovery without compromising peak capacity.

A suggested strategy to optimize chromatography for HDX MS is as follows, based on ideas for peptide separations described by [19, 32] and adapted to HDX MS. 1. Use packing material with sub 2 μm particles. 2. chose column dimensions compatible with the backpressure limit. A 1×50 mm column works well but 2.1×50 mm or 2.1×30 mm may also provide good performance without exceeding backpressure limits. 3. Choose the working temperature – for HDX MS this usually means 0 °C. 4. Operate close to the maximum flow rate (and therefore backpressure) possible with the chromatographic system available. Again, a 1×50 mm column is usually adequate for most separations, even for complex samples. 5. Based on sample complexity, decide if separation speed is more important than peak capacity, then set the gradient time to the longest time tolerable. For peptides, a 5-35% acetonitrile gradient seems the most universal.

Finally, separation in HDX MS is multi-dimensional. There is chromatographic resolving power, ion mobility resolving power (if the mass spectrometer is so equipped), and m/z resolving power. The final peak capacity is therefore multiplex and much higher than just the chromatographic component. For example, [LC × IMS × MS = total peak capacity] could be crudely estimated by peak capacities such as [50 × 50 × 400 = 1,000,000] in a more ideal case. In a less ideal case, the values will be lower, e.g., [10 × 20 × 400 = 80,000], and if there is no IMS [10 × 400 = 4,000]. Such multiplex power is key to being able to extract deuteration information for peptic peptides from even extremely complex digests with many peptides. Even small improvements in any of the three separation mechanisms (LC, IMS, MS) can translate to vast improvements in HDX MS data quality.

## Supporting information

Supplemental Datafile

## Acknowledgements

We gratefully acknowledge the following people for helpful discussions and assistance with this project: Azita Kaffashan, Keith Fadgen, Colette Quinn, Iggy Kass, Lindsay Morrison, Alexandre Gomez, Jim Murphy, Adam Parkinson, and Tom Smith. Financial support was provided by grants R01 CA-233978, R01 AI-043957, R01 GM-067260, R01 GM-135158, and R35 CA-197583 from the National Institutes of Health.

## Supplemental material

Supplemental material associated with this article can be found, in the online version, at doi: XXXXXX.

