## Supplemental Material for "Increase the flow rate and improve hydrogen deuterium exchange mass spectrometry"

Table of contents:

Content description for accompanying *"Supplemental_Datafile.xlsx"* file
A Microsoft Excel file containing: Peak width measurement values, start/end retention times, calculations of sample peak capacity and confidence intervals

Figures S1-S7:
Figure S1. I-Class HDX MS system and additional back pressure measurements
Figure S2. Expanded chromatographic performance information
Figure S3. Details of the values used to calculate the sample peak capacity
Figure S4. Normalized percent change in sample peak capacity
Figure S5. Details of the four-protein mixture
Figure S6. Peak width measurements for the four-protein mixture
Figure S7. Peptic peptides of myoglobin

Supplemental material references

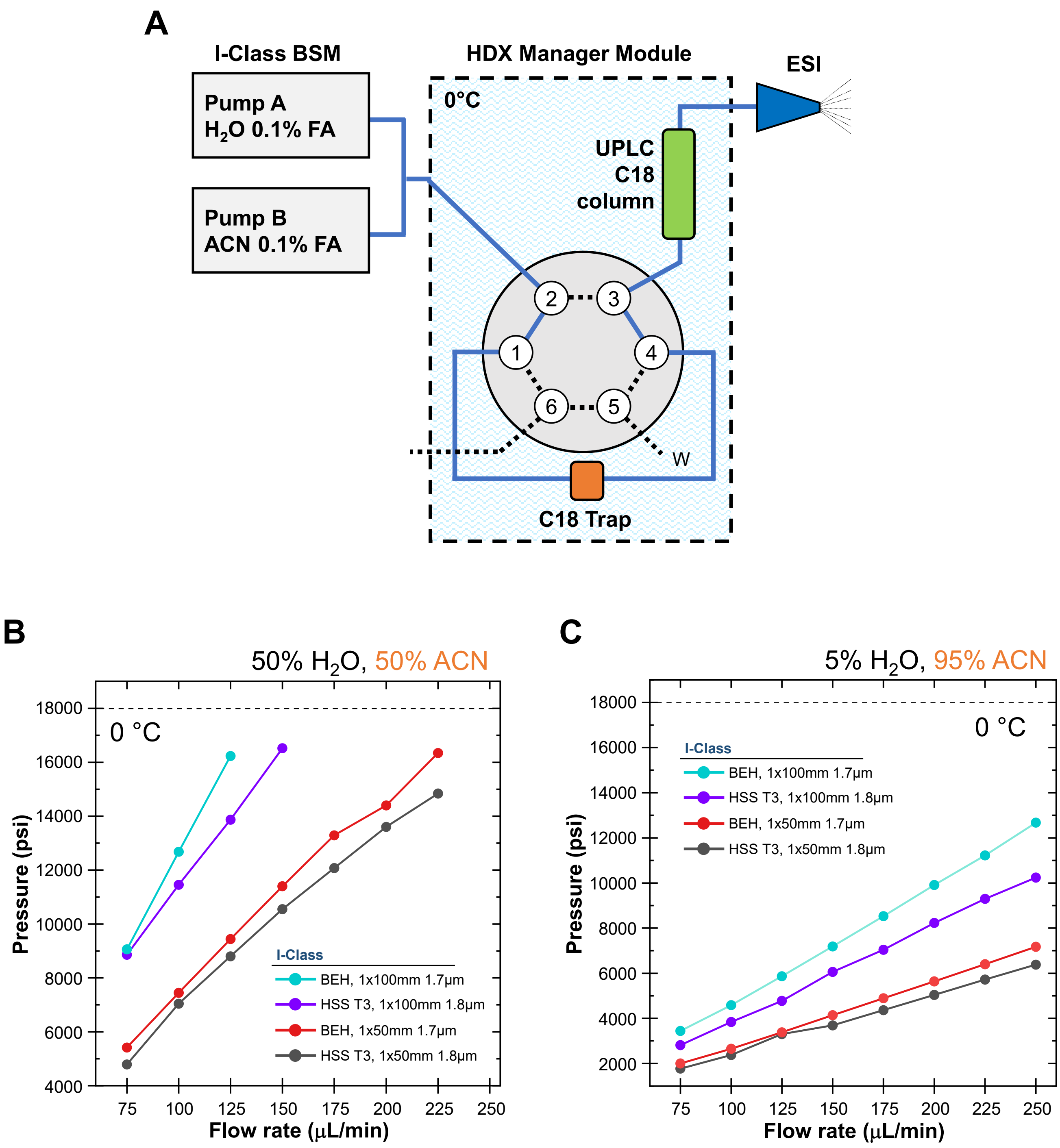


**Figure S1.** I-Class HDX MS system and additional back pressure measurements. (A). Flow path diagram for the I-Class system configured for HDX MS. (B,C). Backpressures measured for the I-Class HDX MS system at 0 °C and solvent compositions (B) 50% acetonitrile 50% acetonitrile, 0.1% formic acid and (C) 95% acetonitrile, 5% water, 0.1% formic acid.

**A**

Corrected gradient steepness ($G_s$)

| Gradient length (min) | Flow rate (µL/min) 40 | 50 | 65 | 75 | 100 | 125 | 150 | 175 | 200 | 225 | 250 |
|---|---|---|---|---|---|---|---|---|---|---|---|
| 3 | 6.68 | 5.34 | 4.11 | 3.56 | 2.67 | 2.14 | 1.78 | 1.53 | 1.34 | 1.19 | 1.07 |
| 6 | 3.34 | 2.67 | 2.05 | 1.78 | 1.34 | 1.07 | 0.89 | 0.76 | 0.67 | 0.59 | 0.53 |
| 10 | 2.00 | 1.60 | 1.23 | 1.07 | 0.80 | 0.64 | 0.53 | 0.46 | 0.40 | 0.36 | 0.32 |

**B**

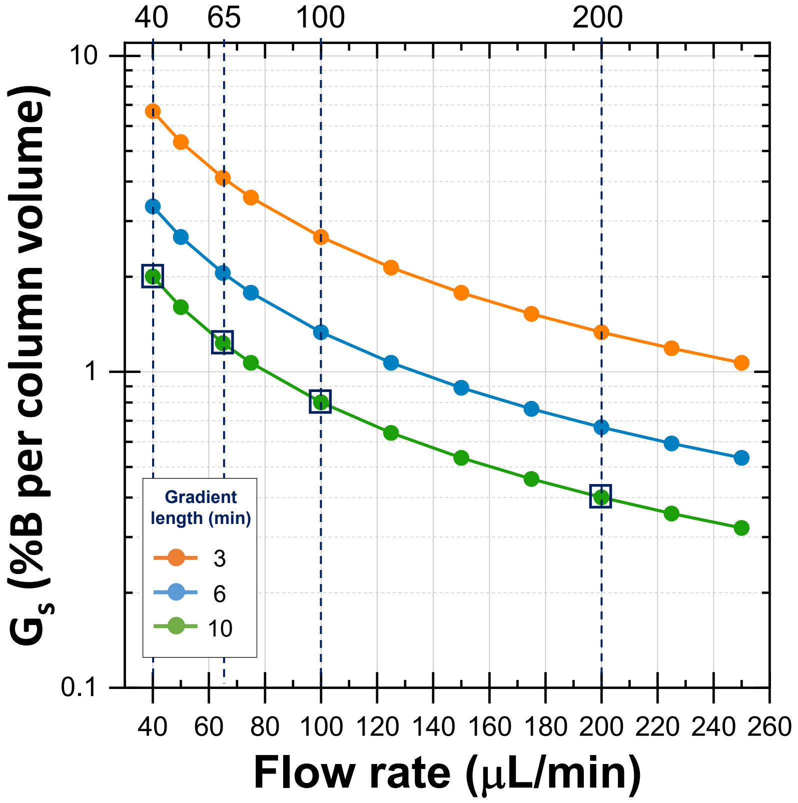


**Figure S2.** Expanded chromatographic performance information. (A). Additional corrected gradient steepness ($G_s$) calculations. Using the same column geometry as in Figure 2, $G_s$ was calculated for combinations of gradient length and flow rate for the gradient lengths in both Figure 2 (3 mins, 6 mins) and Figure 4 (10 mins). This table includes the flow rate values in Figure 2B (40, 75, 100, 150, 200, 225 µL/min) and other useful values (50, 65, 125, 175, 250 µL/min). Dotted boxes highlight the $G_s$ at the flow rate and gradient lengths measured in Figure 4. (B). A graph of the values in panel A, highlighting with dotted lines and boxes the flow rates (40, 65, 100 and 225 µL/min) that were tested in Figure 4.

(Continued…)

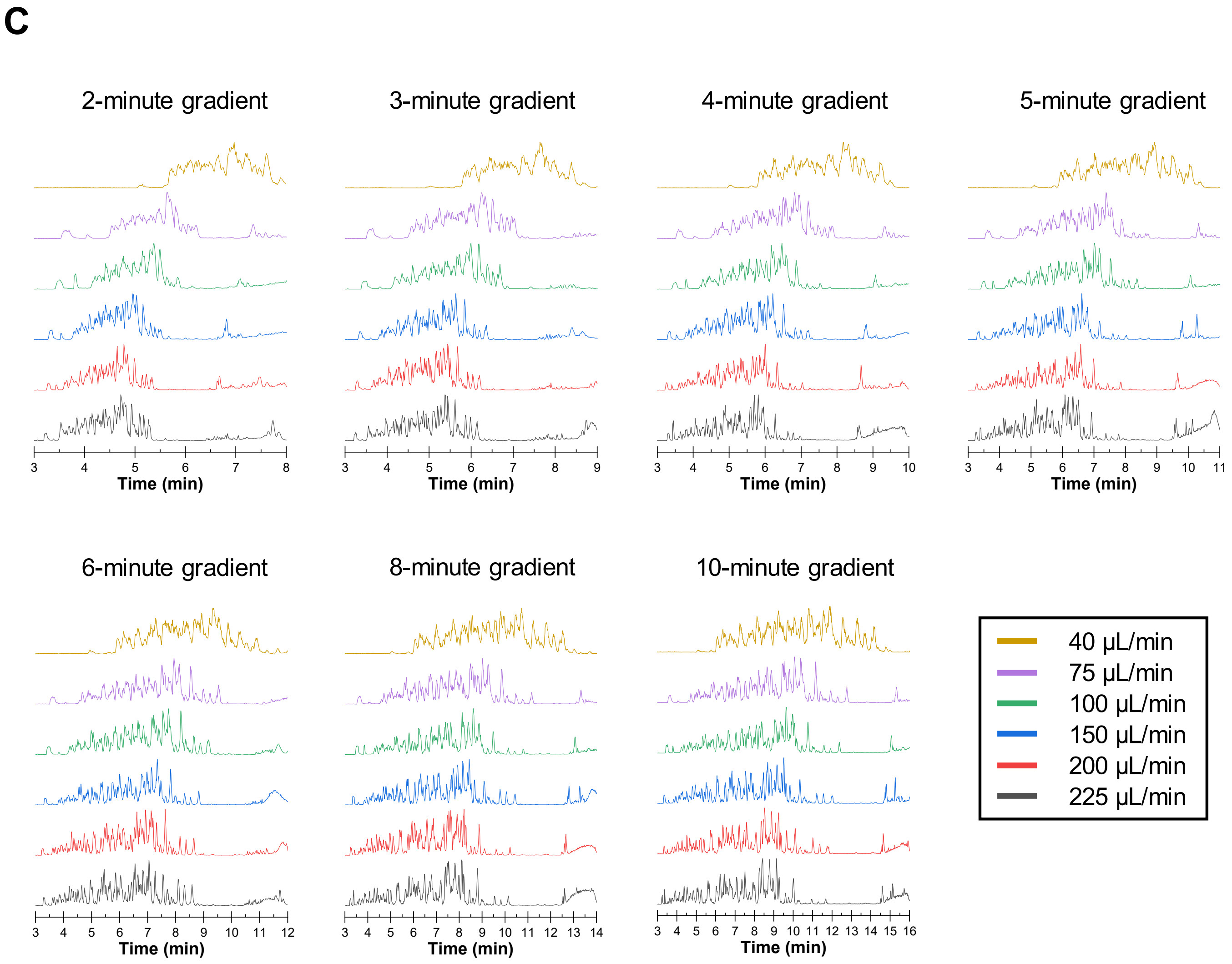


**Figure S2.** (C). Complete set of chromatograms (BPI on the Y axis) of the 0 °C separation of a phosphorylase b tryptic peptide digest (8 pmol injection, MassPREP standard, Waters) on a Waters HSS T3 column (1×50mm, 1.8 µm particles, 100 Å pores). The gradient was 5-35% acetonitrile (in water with 0.1% formic acid) with gradient length and flow rates as indicated. Base-peak intensity (BPI) has been plotted.

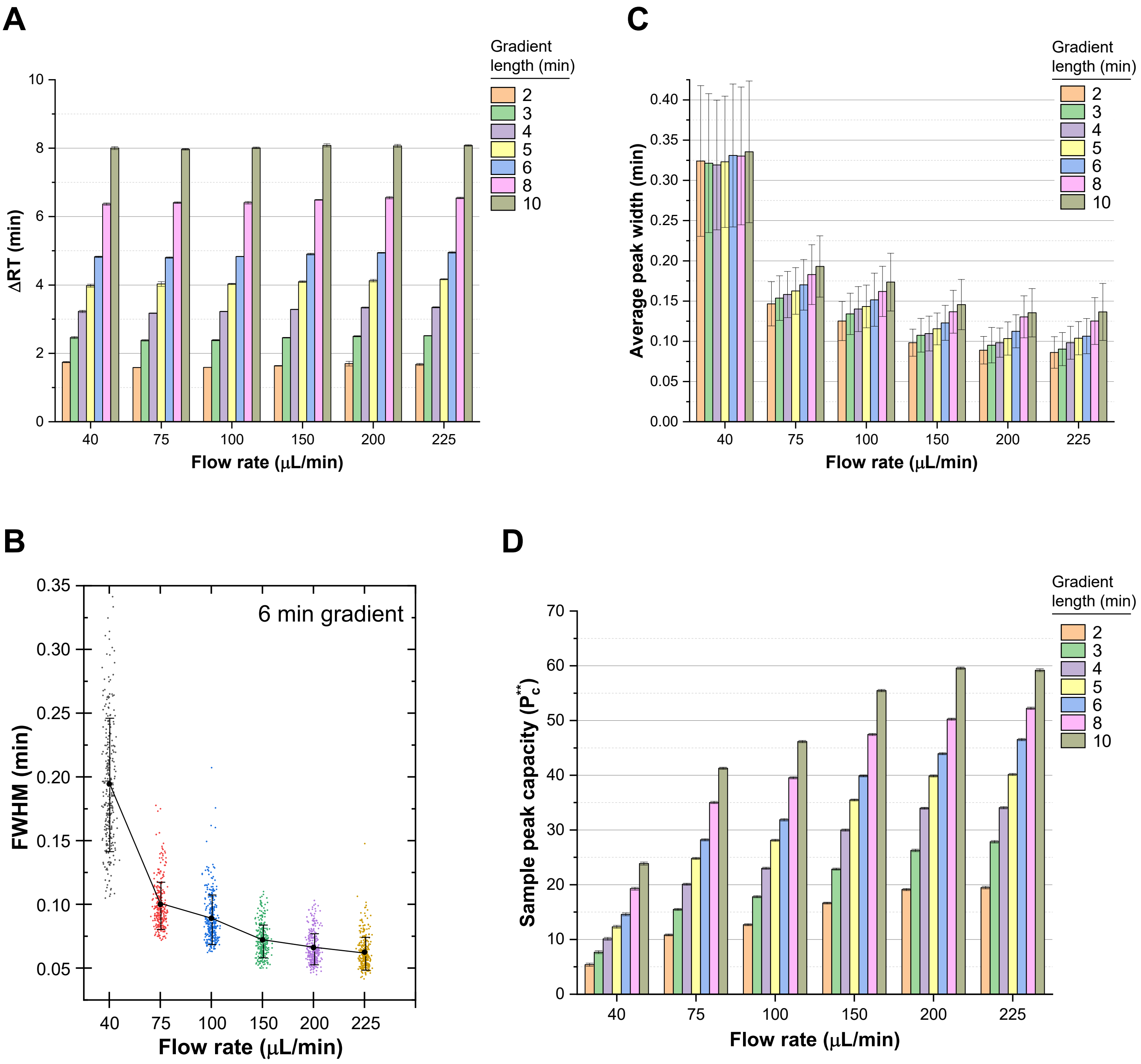


**Figure S3.** Details of the values used to calculate peak capacity. All values, including error values, can be found in the Supplementary Datafile. (A). ΔRT versus flow rate for each gradient length, where ΔRT is the difference in retention time between the last ($t_l$) and first ($t_f$) eluting species, i.e., the top portion of Eq. 2. The error bars represent the (B). Distribution of peak widths, shown (for example) for the 6 minute gradient data. PLGS was used to measure the full width half maximum (FWHM) peak width for the 100 most intense peaks/peptides in each separation at each flow rate, in triplicate. All 300 peak widths (dots) from the triplicate measurements are plotted, along with the average peak width (at FWHM) of all 300 measurements, and the (error bars) average standard deviation (at FWHM) of all 300 measurements. Measured values for other retention times are in the Supplementary Datafile. (C). Average peak width at 4σ versus flow rate for each gradient length. The error bars are average standard deviation at 4σ. (D). Sample peak capacity ($P_c^{**}$) versus flow rate for each gradient length. These values are the same as those shown in Figure 3, error bars represent the confidence interval at 95% confidence.

**A**

Normalized % change, sample peak capacity ($P_c^{**}$)

Flow rate (µL/min)

| Gradient length (min) | 40 | 75 | 100 | 150 | 200 | 225 |
|---|---|---|---|---|---|---|
| 10 | -25% | 29% | 45% | 74% | 87% | 86% |
| 8 | -40% | 10% | 24% | 49% | 58% | 64% |
| 6 | -54% | -12% | 0% | 25% | 38% | 46% |
| 5 | -61% | -22% | -12% | 11% | 25% | 26% |
| 4 | -68% | -37% | -28% | -6% | 7% | 7% |
| 3 | -76% | -51% | -44% | -28% | -18% | -13% |
| 2 | -83% | -66% | -60% | -48% | -40% | -39% |

**B**

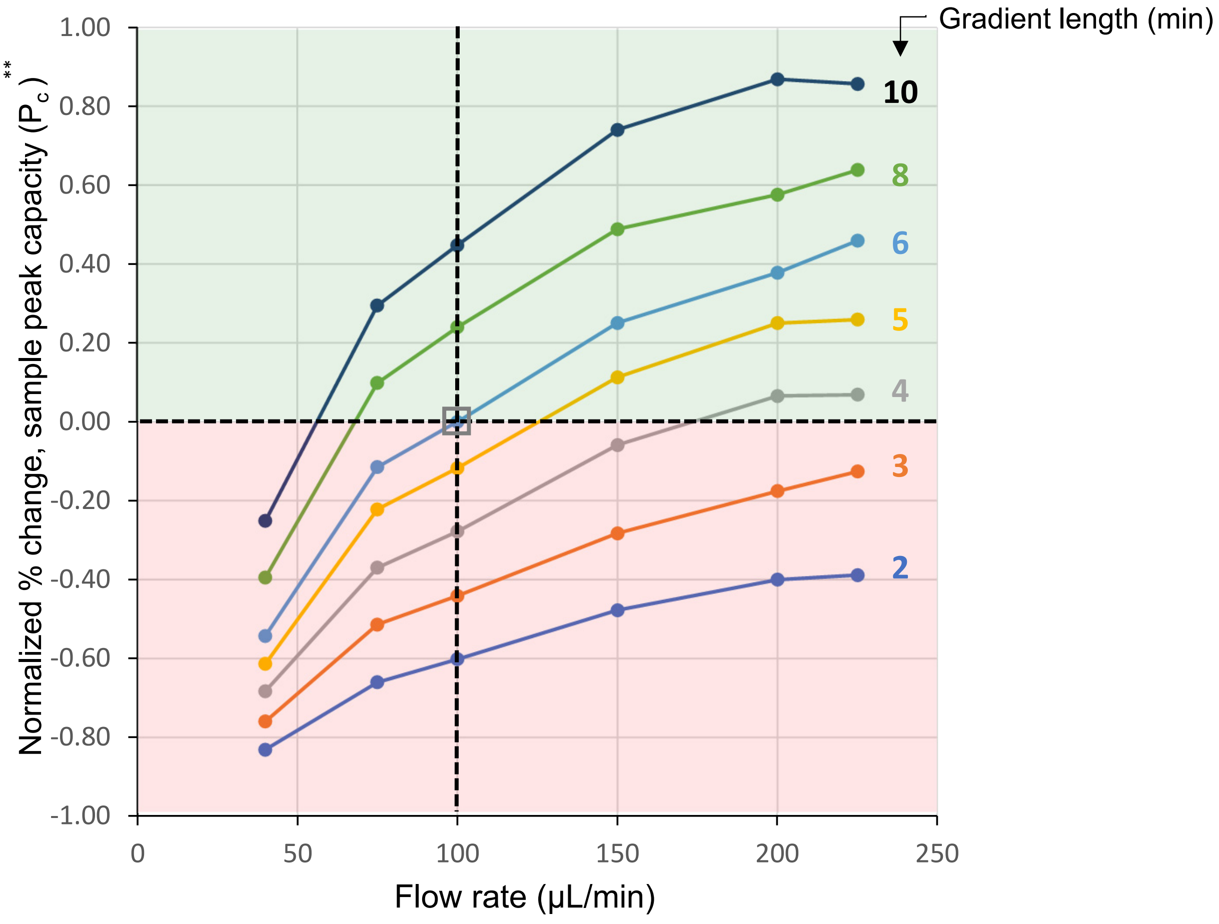


**Figure S4.** Normalized percent change in sample peak capacity. (A). The percent sample peak capacity change was calculated from the $P_c^{**}$values (Figure 3B) determined for each flow rate:gradient length combination according to the formula [($P_c^{**}/P_{c,reference}^{**}$)-1] where $P_{c,reference}^{**}$ was the flow rate and gradient length combination of 100 µL/min and 6 min gradient (dotted box). The resulting percent change values were colored from worst (red) to best (green). (B). A graph of the values from panel A where the reference point is 100 µL/min, 6 min gradient (gray boxed value at center). The normalized percent change values are plotted according to gradient length (left side). Positive or negative change is highlighted by a green or red shaded region, respectively.

| | ADH | Enolase | BSA | PhosB |
|---|---|---|---|---|
| Full name | Alcohol dehydrogenase | Enolase | Serum albumin | Glycogen phosphorylase |
| Species | *S. cerevisiae* (yeast) | *S. cerevisiae* (yeast) | *B. taurus* (cow) | *O. cuniculus* (rabbit) |
| Monomer residues | 347 | 436 | 583 | 842 |
| Average MW monomer (Da) | 36,718 | 46,685 | 66,433 | 97,158 |
| Quaternary structure | Homotetramer | Homodimer | Monomer | Homodimer |
| Average MW quaternary (Da) | 146,872 | 93,370 | - | 194,316 |
| Sigma part # | A3263 | E6126 | A8531 | P6635 |
| Quantity monomer injected (pmoles) | 20 | 20 | 20 | 20 |
| Uniprot Code | P00330 | P00924 | P02769 | P00489 |
| Example PDB Code | 4W6Z | 1EBG | 6QS9 | 1A8I |
| | 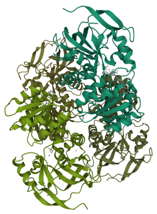 | 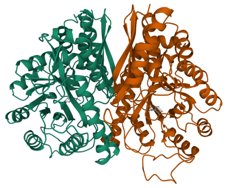 | 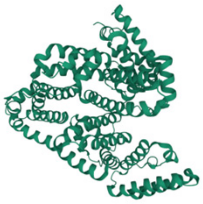 | 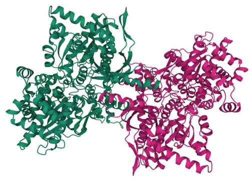 |

Mixture properties: **501 kDa Mass quantity**

**2,208 Unique sequence**

**Figure S5.** Details of the four-protein mixture. The mixture, as described previously (Wales, Fadgen et al. 2008), was prepared from the components shown and had a mass quantity (MQ) of ~501 kDa and 2,208 amino acids of unique sequence (UqSeq). MQ is the sum of the molecular weights of the biologically occurring multimeric forms (here MQ: 500,991 = 146,872 + 93,370 + 66,433 + 194,316), UqSeq is the sum of the number of residues from each monomer with unique sequence (here UqSeq: 2,208 = 347 + 436 + 583 + 842). These terms are fully explained in (Engen and Komives 2020).

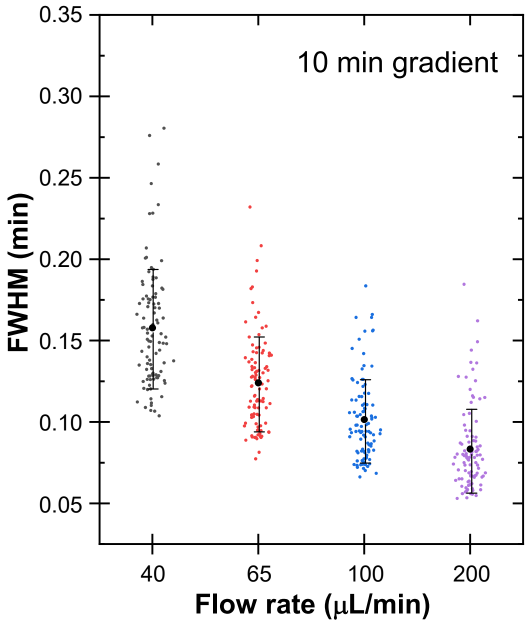


| Flow rate (μL/min) | Average FWHM (min) | StdDev. |
|---|---|---|
| 40 | 0.158 | 0.037 |
| 65 | 0.124 | 0.030 |
| 100 | 0.102 | 0.026 |
| 200 | 0.083 | 0.026 |

**Figure S6.** Distribution of peak widths in the four-protein mixture chromatograms. PLGS was used to measure the full width half maximum (FWHM) peak width for the 100 most intense peaks/peptides in each separation at each flow rate, in triplicate. 100 measurements (dots) from the first replicate are plotted as an example, along with the average peak width (at FWHM), and the (error bars) average standard deviation (at FWHM). The average values and standard deviations from 40 to 200 μL/min are shown in the table.

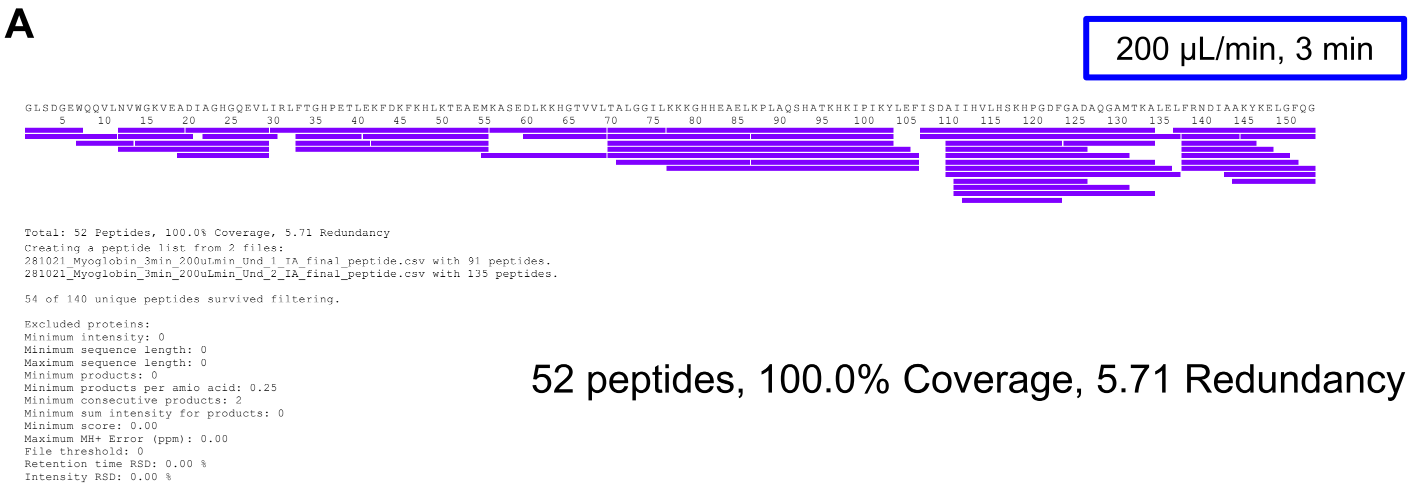


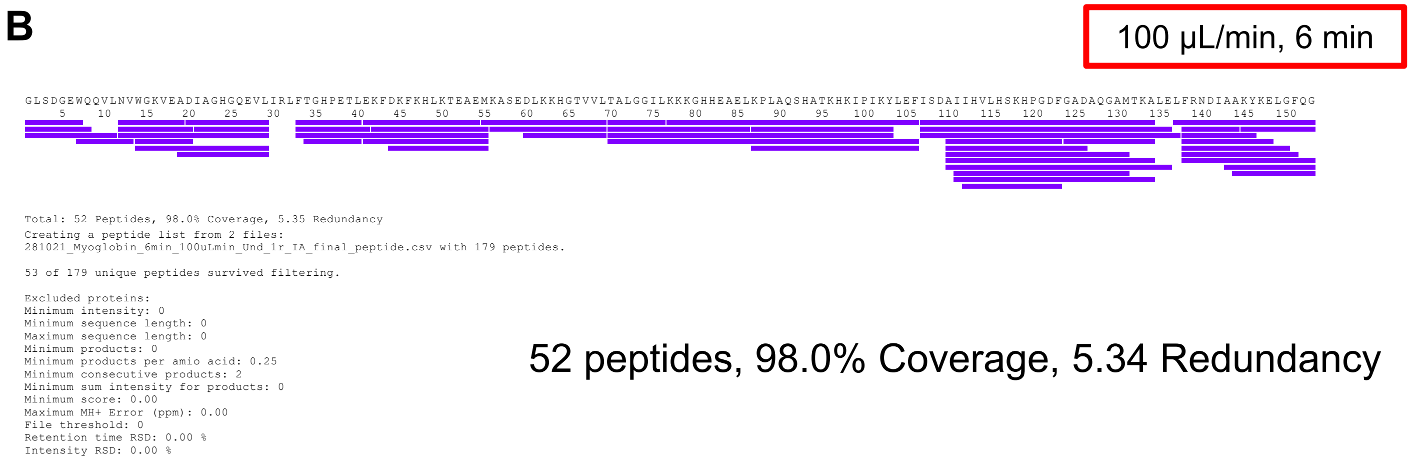


**Figure S7.** Peptic peptides of myoglobin identified in Figure 5. Nearly the same peptides were seen at both flow rate:gradient combinations, (A) 200 µL/min, 3 min gradient and (B) 100 µL/min, 6 min gradient.

### Supplemental material references


Engen, J. R. and Komives, E. A. (2020). Complementarity of Hydrogen/Deuterium Exchange Mass Spectrometry and Cryo-Electron Microscopy. ***Trends Biochem Sci*** **45**(10): 906-918.

Wales, T. E., Fadgen, K. E., Gerhardt, G. C. and Engen, J. R. (2008). High-speed and high-resolution UPLC separation at zero degrees Celsius. ***Anal Chem*** **80**(17): 6815-6820.
